# Control of Astrocyte Quiescence and Activation in a Synthetic Brain Hydrogel

**DOI:** 10.1101/785683

**Authors:** Sualyneth Galarza, Alfred J. Crosby, ChangHui Pak, Shelly R. Peyton

## Abstract

Bioengineers designed numerous instructive brain extracellular matrix (ECM) environments that have tailored and tunable protein composition and biomechanics *in vitro* to study astrocyte reactivity during trauma and inflammation. However, a major limitation of both protein-based and model microenvironments is that astrocytes within fail to retain their characteristic stellate morphology and quiescent state without becoming activated under “normal” culture conditions. Here we introduce a synthetic hydrogel, that for the first time demonstrates maintenance of astrocyte quiescence, and control over activation on demand. With this synthetic brain hydrogel, we show the brain-specific integrin-binding and matrix metalloprotease (MMP)-degradable domains of proteins control astrocyte star-shaped morphologies, and we can achieve an ECM condition that maintains astrocyte quiescence with minimal activation. In addition, we can induce activation in a dose-dependent manner via both defined cytokine cocktails and low molecular weight hyaluronic acid. We envision this synthetic brain hydrogel as a new tool to study the physiological role of astrocytes in health and disease.

---

Astrocytes constitute approximately 30%^1^ of the cells within the mammalian brain and function as key producers and maintainers of the brain extracellular matrix (ECM) during brain tissue homeostasis.^2,3^ During brain trauma^4^ and inflammation^1,5^, changes in the ECM composition^6-10^, ECM stiffness^11^, and the introduction of cytokine molecules^12^ transform astrocytes from a quiescent to a reactive state. This reactive state is typically characterized by the upregulation of the intermediate filament proteins glial fibrillary acidic protein (GFAP)^1-3^ and vimentin^13^. Recent studies^1^ have sought to understand the profile and origination of reactive and quiescent astrocytes^12,14^ toward developing therapeutics that inhibit astrocyte activation^1^. Yet, these studies are hindered due to the large complexity and lack of control over the *in vivo* astrocyte microenvironment. Thus, researchers have increasingly sought *in vitro* models in which to study astrocyte activation. However, a major limitation to the study of normal and healthy astrocytes *in vitro* is that there is no reproducible system that can maintain astrocytes in a quiescent state to study activation.

For this reason, bioengineers have developed defined and controllable cell culture environments in which to study cells in more native-like conditions. Relevant to the brain, astrocytes grown as three-dimensional (3D) organoids^15,16^ provide aspects of their native phenotype, such as a stellate morphology and astrocyte cell-to-cell heterogeneity. However, the close cell-cell contact in these organoid culture is an important drawback, as astrocyte processes only overlap *in vivo* during their reactive state.^2,17^ Additionally, this culturing cells as organoids is time consuming^15,18^ and does not allow for the customization of ECM cues like stiffness^6,19,20^ and ligand density^7,20^ present in real tissue. Protein and glycosaminoglycan-based 3D hydrogels, *e*.*g*. Type I collagen^7,9^, hyaluronic acid (HA)^21^, or defined mixtures of the two^7,9^ have been popularized as *in vitro* models of brain ECM as they are naturally occurring in the brain, are biocompatible, and have shown lower upregulation of GFAP^9^ compared to astrocytes grown as a 2D monolayer. These environments still cause significant astrocyte activation compared to *in* vivo, and a major drawback of naturally occurring ECM model systems is that their constituent proteins^6,8-10^, and stiffnesses^6^ (which likely influence astrocyte activation) cannot be independently controlled.

Synthetic hydrogels, in contrast, provide a tremendous opportunity to design tissue-specific scaffolds^20^ with tight control over environmental parameters, and there is an engineering opportunity to incorporate bioactive molecules to represent any microenvironment of interest.^22-24^ However, astrocytes either become reactive when cultured in synthetic hydrogels^23^, or in cases where activation was reduced, their characteristic stellate morphology was sacrificed^25,26^. Currently there is no tunable *in vitro* model that can maintain and control astrocyte quiescence, and therefore further the study of how astrocytes activate during central nervous system (CNS) diseases or other injuries/trauma. This highlights the imminent need for ECM environments that can retain astrocyte physiological quiescence *in vitro*.

We sought to develop such an *in vitro* model that would enable us and others to study the specific extracellular factors that control astrocyte quiescence and activation. To do this, we first characterized the human brain ECM via mechanical indentation, mass spectrometry, and Protein Atlas histology^27^ in order to incorporate the appropriate stiffness and most prevalent ECM proteins responsible for integrin-mediated adhesion and MMP-mediated degradation in the brain. We synthesized peptides to represent these proteins, and combined them with a modulus-tunable poly(ethylene glycol) (PEG) network to create a carefully designed, yet very simple synthetic hydrogel brain-like ECM. With this hydrogel we overcome the current challenge of retaining primary human astrocyte quiescence *in vitro* and, therefore, controlling activation. We demonstrate control over astrocyte reactivity via tuning of the integrin-binding and MMP-degradable profile of the hydrogel, or via dosing with cytokine molecules and low molecular weight HA, currently not possible in other *in vitro* systems where astrocytes remain in a permanently activated state.

## Characterization of the Human Brain ECM

Astrocytes are responsive to ECM proteins *in vitro*. For example, ECM composition^6^ determines astrocyte responses to substrate stiffness and other inflammatory stimuli. These authors observed arrested astrocyte migration on fibronectin, and rapid migration on tenascin or laminin. Others have shown astrocyte attachment to fibronectin, laminin, and fibrillin-1 regulates IL-1-induced activation in an integrin-dependent manner.^28^ Integrin heterodimers are the transmembrane receptors that mediate tissue-specific cell binding to the ECM. We thus hypothesized that a brain-specific ECM with defined integrin-binding interactions would allow for control over astrocyte quiescence and activation *in vitro*. To define the ECM components of real brain tissue, we acquired four healthy human frontal cortex samples (Fig. 1a). These samples were decellularized and enriched for ECM proteins following an ECM enrichment protocol recently introduced by Naba *et al*,^29^ which resulted in an insoluble pellet of 1-2 weight % ECM, which we then analyzed via Liquid-Chromatography Mass-spectrometry (LC-MS/MS) (Fig. 1a). Protein hits were compared to the Human Matrisome Database^30^ and classified into ECM-core or -related proteins (Fig. 1b). Glycoproteins, collagens, and proteoglycans, which are known to interact with integrin heterodimers, compose the core ECM proteins (Fig. 1b). From this list, and limiting to hits found in at least two donors (Fig. 1c), proteins were further stratified by affinity to integrin binding and enzymatic degradation via matrix metalloproteinases (MMPs, Fig. 1d-e, Table S2). Peptide spectrum match (PSM) normalized by the protein molecular weight was used to quantify a relative abundance of selected proteins (Fig. 1d-e). At the culmination of this process, we found 116 ECM proteins, 18 of which with known integrin interactions, and 21 known to be degraded by MMPs (Fig. 1d-e, Table S1-2).

**Figure 1.**
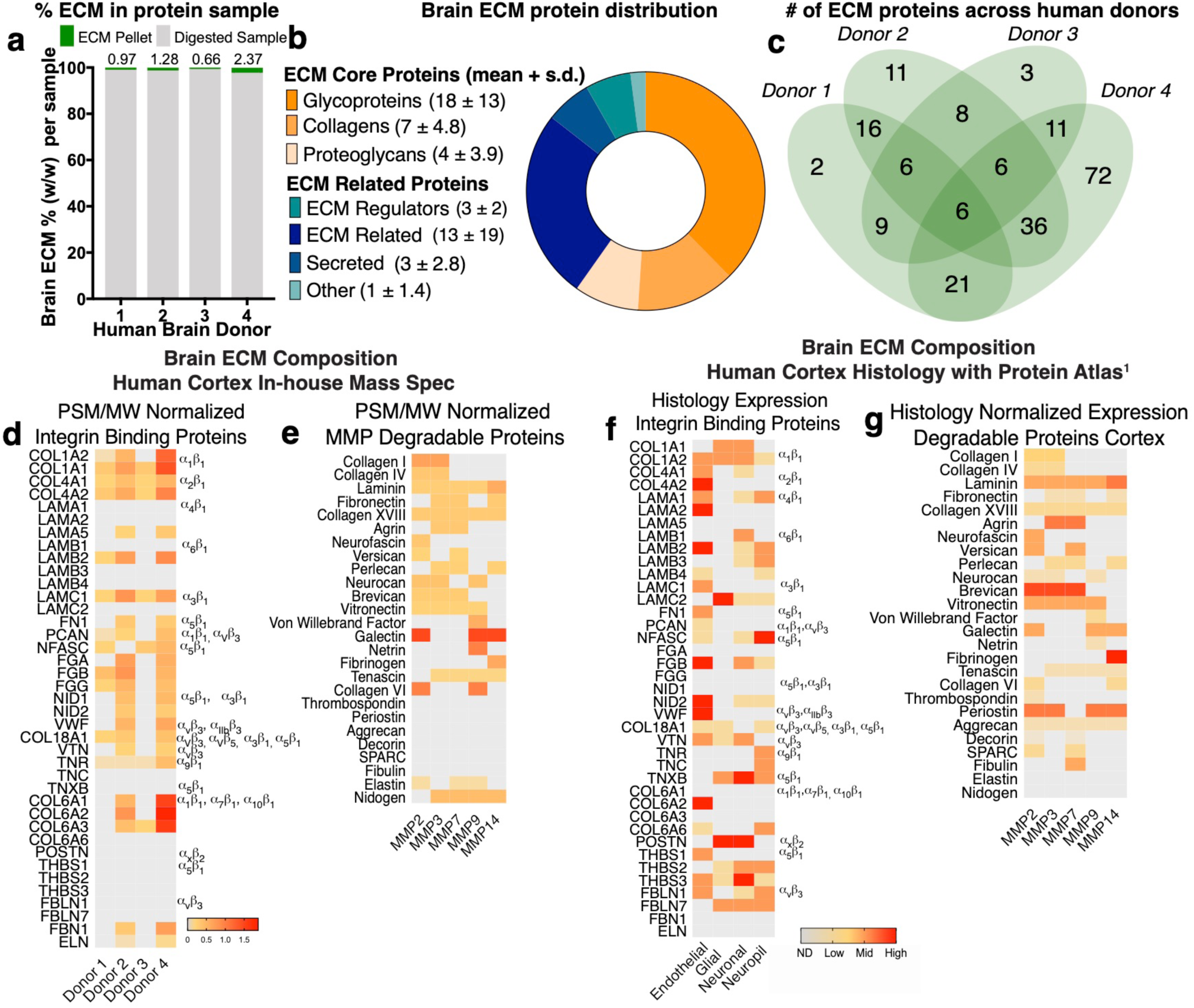
Characterization of the Human Brain Cortex Extracellular Matrix. **a** Right frontal cortex samples from four healthy human donors were decellularized and enriched for ECM proteins - resulting in an insoluble pellet that was solubilized, reduced, and digested into peptides identified via Liquid Chromatography-Mass Spectrometry (LC-MS). **b**. Protein hits were compared to the Human Matrisome Database and classified into ECM-core or -related proteins. Data shown is average number of proteins found across the four donors + standard deviation (s.d.). **c**. Distribution of brain ECM signature proteins among donors **d**. Proteins identified in at least two donors were screened for affinity to integrin binding and **e**. enzymatic degradation via matrix metalloproteinases (MMP). Heat map depicts protein abundance based on peptide spectrum match (PSM) normalized to protein molecular weight (MW) for Integrin binding- and MMP-degradable proteins. **f**. The Human Protein Atlas was screened for ECM proteins in the human brain cortex as identified via the provided histology. Heat map depicts protein abundance via histology expression of integrin binding and **g**. MMP-degradable proteins in the human brain cortex. Degree of expression denoted as ND (not detected), low, medium, or high.

To ensure that the proteins we identified were in areas of the brain both near astrocytes and in the extracellular space (not intracellular), we complemented our LC-MS/MS findings with a published histological screen of brain proteins by the Human Protein Atlas^27^, where we found 147 brain ECM proteins in the cortex (Table S3). We similarly reduced this list to core proteins known from the Matrisome database, resulting in 17 proteins with known integrin binding sites and 24 that are degraded by MMPs (Fig. 1f-g, Table S4). Spatial location of proteins within the brain cortex was assessed via histology images from the Protein Atlas to differentiate proteins located at the basement membrane (endothelial cells) or the interstitium (astrocytes, neurons, neuropils, Fig. 1f-g)^31^. We used the grade of histology reported by the Protein Atlas (ND, low, medium, or high) to assess protein abundance (Table S3-4). Both approaches, in-house mass spectrometry and Protein Atlas histology, were combined to determine the relative amounts of proteins present in the brain cortex (Fig. 2a; Table S5).

**Figure 2.**
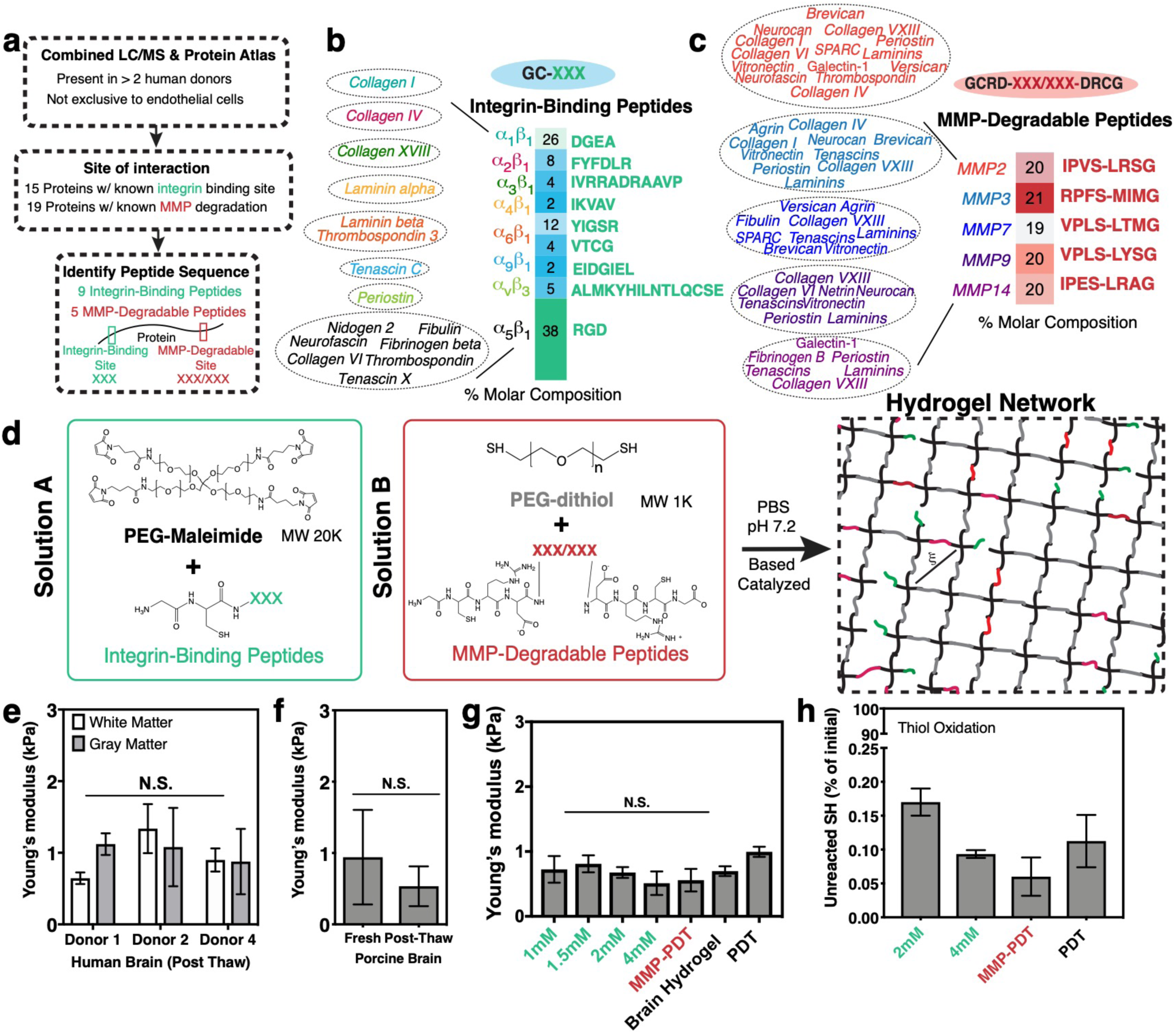
Design of a Synthetic Brain ECM Hydrogel. **a**. Proteins identified via *in-house* proteomics (LC/MS, present in >2 donors) and histology from the Protein Atlas (to exclude proteins specific to endothelial cells) were screened by integrin-binding and matrix metalloproteinase (MMP)-degradable peptide sequences. **b**. Integrin-binding peptides include fifteen proteins, incorporated into the hydrogel via single cysteine. Heat map depicts molar concentration of each peptide in the final design. **c**. MMP-degradable peptides include nineteen proteins, and were incorporated into the hydrogel by a cysteine at each end. Heat map depicts molar concentration of each peptide in the final design. **d**. A Michael-addition reaction is used to combine these peptides with PEG-maleimide to form a hydrogel network via two solutions: solution A containing integrin-binding peptides and the PEG polymer, and solution B with the MMP-degradable peptides and a non-degradable crosslinker. ξ is the mesh size of the hydrogel (approximately 20nm). **e**. Young’s modulus of previously frozen human brain (3 donors) **f**. fresh and previously frozen porcine brain *n=6* and **g**. hydrogel tuned to the same Young’s modulus as brain tissue as measured via indentation. Modulus is maintained after incorporation of integrin-binding (green) or MMP-degradable (red) peptides. *n=8*. **h**. Percent of unreacted thiols after hydrogel is formed, then reduced with NaBH_4_, indicating high network formation efficiency. *n=8*. All data are mean +/-s.d. Data in **e-g** was analyzed using a ANOVA followed by a Tukey’s multiple comparison test with 95% confidence interval. N.S. is not significant.

## Design of a Synthetic Brain ECM Hydrogel

To create a brain ECM-mimicking hydrogel, we narrowed the ECM Matrisome quantification results to proteins with known interactions with integrin heterodimers.^32,33^ From the pool of proteins present in Figs. 1d and f, integrin-binding sequences were matched via literature mining to their integrin heterodimer pairs (Tables S2,S4,S5). Proteins identified were screened to those present in at least two donors, not present only near endothelial cells, and with known integrin-binding amino acid sequences (Figs. 2a-b). Seven of these fifteen proteins interact with integrins via the RGD site while eight proteins had unique amino acid sequences – resulting in a total of 9 integrin-binding peptides that represent the integrin-binding ECM protein landscape of the synthetic brain ECM (Figs. 2a-b).

Matrix metalloproteinases (MMP) are a family of enzymes that cleave ECM proteins.^34^ Since the PEG-maleimide synthetic hydrogel system has mesh sizes on the order of 10s of nanometers^35^, cells must degrade the biomaterial in order to extend processes, proliferate, and migrate^36^. We screened brain ECM proteins with the same criteria established for integrins as described above (Fig. 2a) and identified those susceptible to MMP degradation. These were grouped based on the MMP via literature mining (19 proteins, Table S2 and S4, Fig. 2a and 2c). Within this group, we identified the specific peptide sequences known to be degraded by MMPs (Figs. 2a and 2c)^37^. This process resulted in 5-degradable peptides that comprise the degradable landscape of the synthetic brain ECM (Fig. 2c).

Both integrin-binding and MMP-degradable peptides can be easily incorporated in the hydrogel during crosslinking via the Michael addition reaction.^38^ Integrin-binding peptides contain a single cysteine functionalization, and MMP-degradable peptides contain cysteines near each end to serve as degradable crosslinks (Fig. 2d). We form the hydrogel by creating two solutions that contain a 1) maleimide functionalized PEG with the integrin-binding peptides, and 2) a PEG-dithiol with the MMP-degradable peptides (Fig. 2d). When the two solutions are combined, the hydrogel network forms in under a minute. We previously showed that this reaction speed can be tuned by manipulating the ionic strength of the buffer, the pH, and the electronegativity of the amino acids flanking the MMP-degradable peptides.^39^

Since brain cell phenotype is sensitive to the modulus of the ECM^40-43^, we needed to account for this in our design of an *in vitro* brain ECM. Unfortunately, we quickly discovered that reported values for Young’s moduli of the brain ranged from 100s of Pa to 10s of kPa, and vary considerably among experimental techniques.^44^ We therefore independently characterized the modulus of human brain, focusing on the cortex, via an indentation technique previously published by our group^45^, and we found the Young’s modulus to be approximately 1kPa (Fig. 2e). Due to our inability to acquire fresh human brain tissue, and to confirm there was no difference in modulus of fresh tissue vs. frozen, we also characterized the modulus of fresh and frozen-then-thawed porcine brain cortex.^46^ We were surprised to find a similar Young’s modulus when comparing porcine and human brain, and no statistically significant difference between fresh and previously frozen porcine tissue (Figs. 2e-f). To incorporate this modulus into our brain hydrogel, and taking a similar approach to work by others^47^ that have modulated a biomaterial’s Young’s modulus, we adjusted the macromer concentration until reaching a similar Young’s modulus to that of brain (Fig. S1). We also confirmed that the Young’s modulus was not affected by the incorporation of a range of concentrations of integrin-binding peptides, or by crosslinking the gel with either PEG-dithiol (PDT) vs. the desired combination of PDT and the MMP-degradable peptides (Fig. 2g). Finally, these peptides can be incorporated efficiently into the PEG-maleimide system as confirmed by an Ellman’s assay and LC/MS (Fig. 2h, Fig. S1).

## Human Brain-Specific Integrin-Binding Peptides Validation via Cell Adhesion

To validate the function of the integrin-binding peptides in the brain-specific hydrogel, we performed a 2D adhesion assay with a human neuroblastoma cell line SK-N-AS. Neuroblastoma cell area increased significantly when cells were seeded on surfaces functionalized with integrin-binding peptides compared to negative control surfaces (PEG, Figs. 3a-b, Fig. S2). Experiments with the breast cancer cell line MDA-MB-231 showed similar results (Fig. S2, Tables S1-4).^48,49^ We conversely performed a competitive binding assay where cells were pre-incubated with soluble, individual integrin-binding peptides before seeding onto a surface with the full cocktail of brain-specific integrin-binding peptides (Fig. 3c). Compared to those cells pre-incubated with peptide, control cells fully spread and adhered to the brain peptide surfaces (Fig. 3d, Fig. S2). We used Pubmed to compare the sequence homology of these peptides found in human ECM proteins to murine (Fig. S3a-i). Murine cell lines Neuro2A and mHippoE18 did not adhere as strongly to the human peptide sequences (Fig. S3c-d), and the competitive binding assay did not inhibit mouse cell adhesion to coverslips, particularly in comparison to the human SK-N-AS cells (Fig. S3g-h).

**Figure 3.**
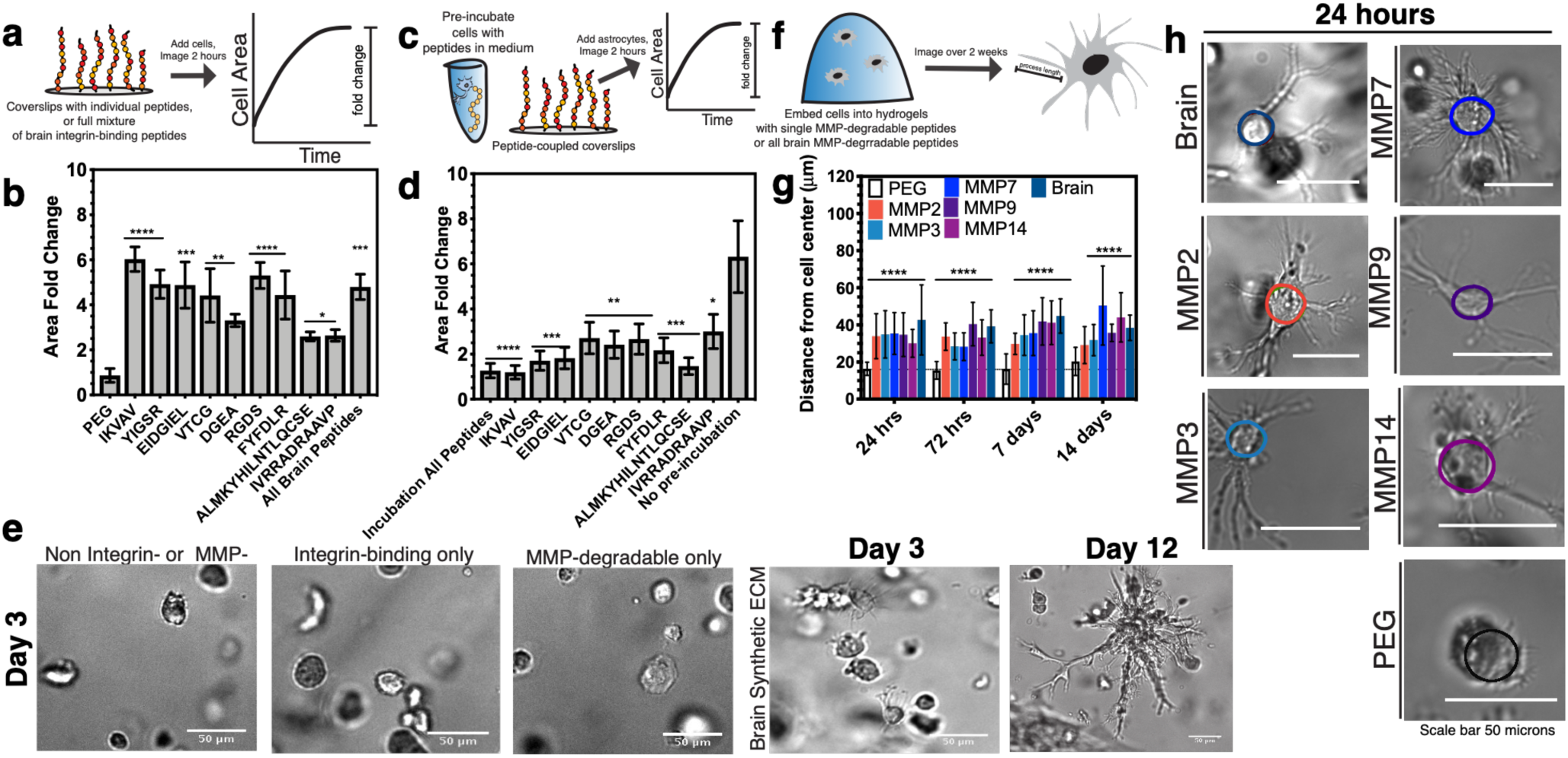
Biological validation of brain integrin-binding and MMP-degradable peptides. **a**. Schematic depicting coverslip functionalization with integrin-binding peptides and cell seeding. **b**. Cell area fold change after seeding onto integrin-binding functionalized coverslips in comparison to a negative control (PEG). N=2, *n=3*. **c**. Schematic of competitive binding assay of integrin-binding peptides. **d**. Change in cell area after pre-incubation with peptides in solution, before seeding onto peptide-functionalized coverslips. Cell area is depicted as a fold change compared to a positive control (no pre-incubation). N=2, *n=3*. **e**. Representative human primary astrocyte morphology in the different hydrogel conditions. **f**. Schematic depicting encapsulation of cells in 3D hydrogels functionalized with MMP-degradable peptides and measurement of process length originating from the cell center. **g**. Astrocyte process length for individual and all brain MMP-degradable peptides in comparison to a PEG gel with no degradable crosslinks after 24, 72 hours, 7 days and 14 days of encapsulation N=2, *n=4*. **h**. Representative images of astrocytes after 24 hours of encapsulation. All data are mean + s.d. Statistical analyses were performed using Prism (GraphPad). Data in **b, d**, and **f** was analyzed using a one-way analysis of variance followed by a Dunnett’s multiple comparison test with 95% confidence interval. *, **, ***, and **** indicate P<0.05, P<0.01, P<0.001 and P<0.0001.

We then encapsulated human primary astrocytes in hydrogels containing a single integrin-binding peptide for 5 minutes prior to fixing and staining for the integrin heterodimers receptors expected to bind the peptide ligand, based on an assay developed by Li *et al*. (Table S5)^50^. This assay only stains integrins at the cell membrane, and positive staining correlates with ECM binding. We positively identified integrin heterodimers via immunofluorescence for the predicted integrin ligand-receptor pairs in contrast to culture in PEG alone (Fig. S4). Together, this data illustrates that the integrin-binding peptides are human brain specific, and appropriate for incorporation into the hydrogel.

## Human Astrocytes Must Cleave MMP-degradable Peptides to Achieve Star-like Morphologies

In the brain, astrocytes are visually phenotypically characterized by extensive ramification and star-like morphologies^3^. Without MMP-degradable crosslinks, hydrogels of a 20-25nm mesh size would yield rounded astrocytes with little or no ramification.^2,3^ We encapsulated human primary astrocytes in 3D hydrogels that were either blank (PEG-only, negative control), hydrogels with integrin-binding peptides only, or containing both brain integrin-binding and MMP-degradable peptides (Fig. 3e). We optimized astrocyte cell density in the brain hydrogel to prevent cell clustering over long culture periods (Fig. S5). We found that astrocytes were able to extend processes in conditions that incorporated both classes of peptides, and they remained viable in the brain hydrogel (Fig. 3e). We verified that each degradable peptide was susceptible to degradation by the astrocytes by encapsulating cells and measuring process length compared to PEG alone (Fig. 3h, Fig. S6), which agrees with work by others for other cell types^37^.

## Astrocyte Activation can be Controlled with Hydrogel Composition

Recent work by Placone *et al*.^9^ illustrated that modifying the concentration of components in a protein-based hydrogel composed of Type I Collagen, HA, and Matrigel reduced the expression of GFAP, an astrocyte-specific marker indicative of activation, in primary fetal human astrocytes as compared to culture in 3D collagen alone. Similarly, Hara *et al*.^8^ demonstrated that astrocytes become reactive when cultured in a collagen-coated substrate mediated by the integrin-N-cadherin pathway. We thus hypothesized that tuning the composition of the brain hydrogel would have an impact on astrocyte activation as measured by GFAP expression.

While varying the MMP peptides constant, we varied the integrin-binding peptide concentrations in the brain hydrogel and quantified cell spreading (Fig. S7). We found that increasing the concentration of adhesive peptides beyond a concentration of 2mM resulted in astrocytes with long processes similar to those in a collagen-based gel, but above 3mM yielded astrocytes with high levels of GFAP fluorescence similar to that in collagen (Figs. 4a-b).^9^ We thus decided to test if modifying the concentration of MMP-degradable peptides would also have an effect in activation. We found that combining a 4mM concentration of integrin-binding peptides with a 13mol% concentration of MMP-degradable peptides yielded significantly low activation of the astrocytes compared to collagen (Figs. 4c-d, Figs. S7-8). This result agrees with that of others upon ECM deposition *in vivo*, where, for example, increase in protein secretion is correlated with astrocyte activation.^8,51^ Figure 4e shows representative morphologies and GFAP fluorescence in astrocytes encapsulated across these different conditions. We found a particular brain hydrogel formulation that resulted in both low activation (GFAP expression) while maintaining long, stellate morphologies: a 1kPa stiffness with 13 mol% MMP-degradable peptides and 2mM integrin-binding peptides.

**Figure 4.**
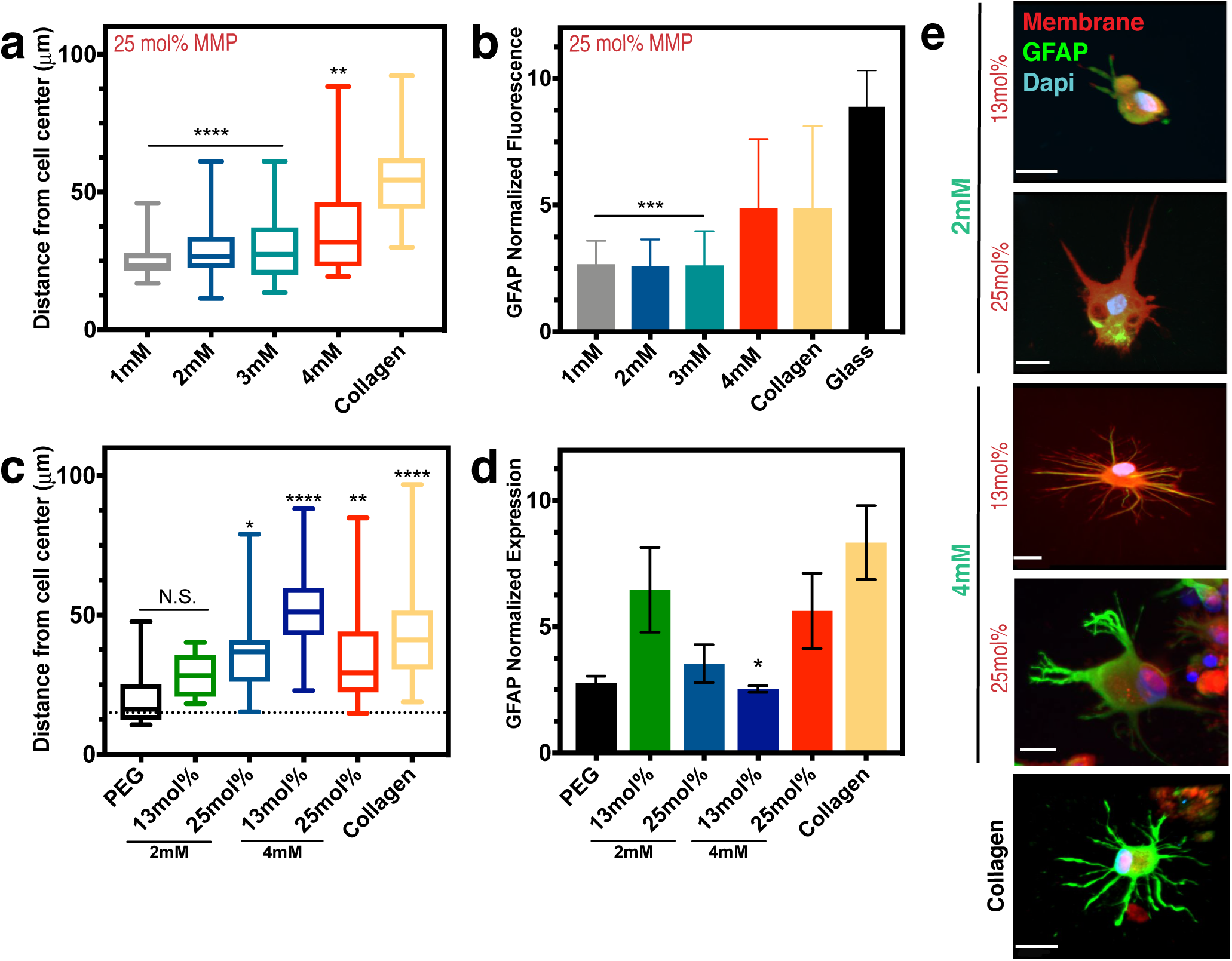
Astrocyte Activation can be Controlled via Integrin-Binding and MMP-Degradable Peptides in the Brain Hydrogel. **a**. Box and whisker plot show distance of process length from the cell center as a function of the integrin-binding peptide concentration (in mM) in the hydrogel, with collagen gels as a comparison. N=2, *n=3*. **b**. Normalized fluorescence intensity of glial fibrillary acidic protein (GFAP) as a function of integrin-specific peptide concentration, with both collagen gels and a glass coverslip for comparison (N =3). All hydrogels in (a) and (b) had 25mol% MMP-degradable peptides and the time point is 48 hours after encapsulation. **c**. Box and whisker plot showing distance of process length from the cell center as a function of both integrin-binding peptide concentration and concentration of the MMP-degradable peptides. Statistics are in comparison to a PEG hydrogel with no peptides included (negative control). N=2, *n=4*. **d**. Normalized GFAP fluorescent intensity as a function of hydrogel integrin-binding and MMP-degradable peptide concentrations. N=2, *n=4*. Data in (c) and (d) is after 72 hours of encapsulation. **e**. Representative images of astrocytes encapsulated in different hydrogel conditions after 72 hours. Scale bar is 20um. Data in (a-c) are mean + s.d. Data in (d) is mean + s.e.m. Data (a-d) were analyzed using a one-way analysis of variance (ANOVA) followed by a Dunnett’s multiple comparison test with 95% confidence interval. *, **, ***, and **** indicate P<0.05, P<0.01, P<0.001 and P<0.0001. N.S. is not significant.

## Astrocyte Quiescence *in vitro* is Maintained in Synthetic Brain ECM Hydrogels

Recent work by Liddelow *et al*.^12^ described astrocyte activation via three cytokines produced by microglia: interleukin 1 alpha (IL-1alpha), tumor necrosis factor alpha (TNF-alpha), and complement component 1, subcomponent q (C1q), resulting in high expression of GFAP.^12^ Similarly, work by Hara *et al*.,^8^ identified *gfap* and *vimentin* as genes highly expressed by spinal cord injury-activated astrocytes. To confirm that the quiescent astrocytes can undergo activation in the brain hydrogel, we dosed them with these three cytokines after 24 hours of culture (Fig. 5a). Under the standard culture medium conditions, astrocytes expressed low levels of activation markers GFAP and Vimentin in the brain hydrogel, in contrast to collagen (Fig. 5b-c).^8^ Similarly, Liddelow *et al*.,^12^ defined a quiescent serum-free medium that is supplemented with heparin-binding EGF-like growth factor (HBEGF). In the brain hydrogel, astrocytes were largely quiescent, but still showed signs of activation in collagen (Figs. 5b-c, Fig. S9). In the brain hydrogel, stimulation with cytokines in increasing doses resulted in increased GFAP expression in the astrocyte population (Fig. 5b-c). In contrast, astrocytes cultured in collagen retained a high GFAP expression across all cytokine concentrations.

**Figure 5.**
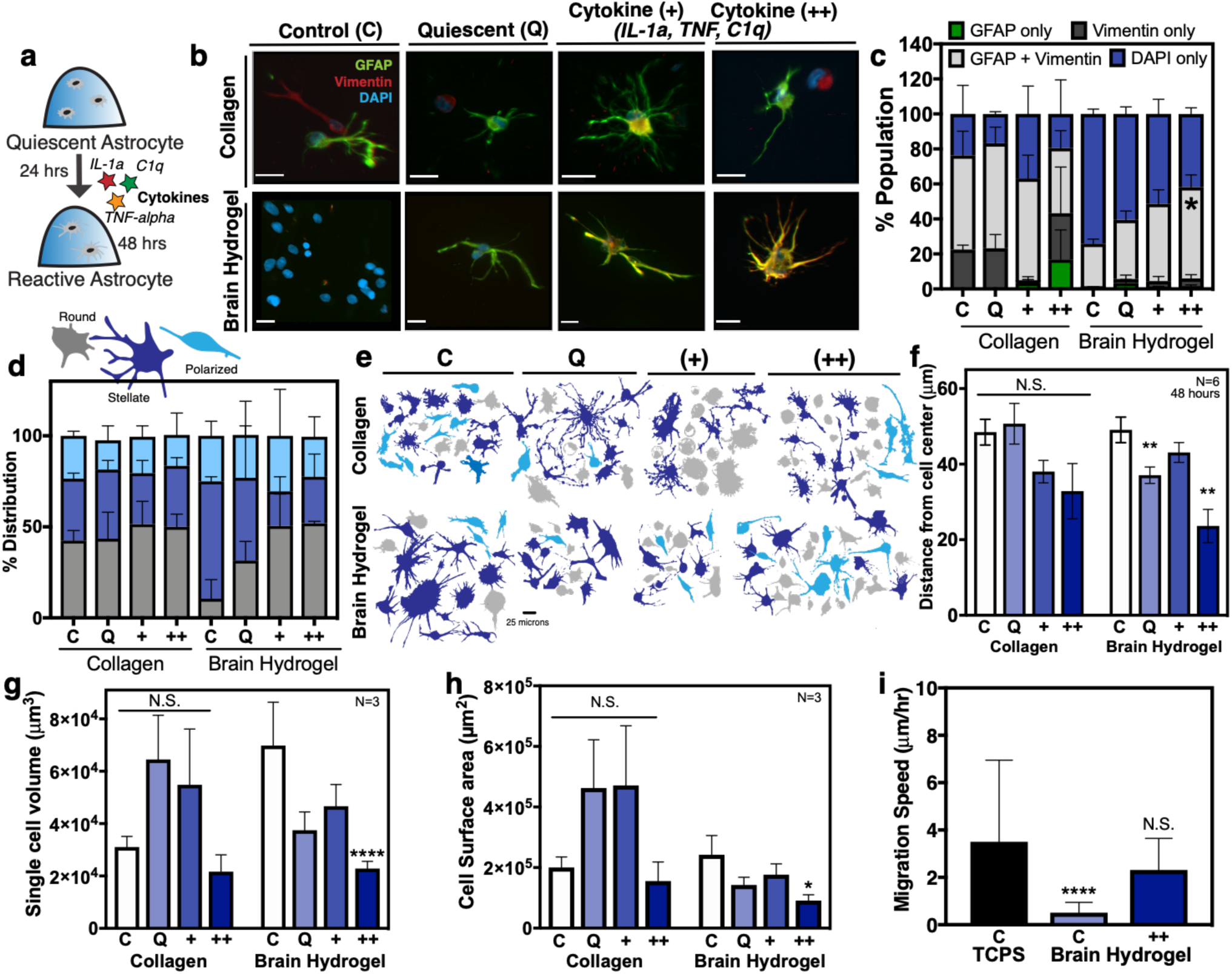
Astrocyte activation can be controlled in the brain hydrogel. **a**. Human primary astrocytes were encapsulated in hydrogels for 24 hours and subsequently dosed with cytokines IL-1*α*, TNF-*α* and C1q for an additional 24 hours. **b**. Representative images of astrocytes after 48 hours in brain hydrogels and collagen and incubated with either standard culture medium (control), defined quiescent medium, dosed with cytokines (+) IL-1*α* (3ng/mL), TNF-*α* (30ng/mL) and C1q (400 ng/mL), or dosed with a 2x dose of cytokines (++) IL-1*α* (6ng/mL), TNF-*α* (60ng/mL), and C1q (800 ng/mL). Scale bar 20 microns. **c**. Quantification of cells from (b) stained for GFAP, vimentin, and DAPI. N=2, *n=250* cells per condition. **d**. Distribution of cell morphologies and **e**. representative cell morphologies in the different populations identified as round (gray), stellate (blue) or polarized (light blue). N=3, *n = 100* cells per condition. **f**. Quantification of astrocyte process length N=6, *n=3*. **g**. Single cell volume and **h**. Single cell surface area for astrocytes encapsulated for 48 hours. N=3, *n=3*. **i**. Comparison of migration speed astrocytes seeded on plastic (TCPS), or in the brain hydrogels with and without cytokines. N=2. All plots show mean + SEM. Data in **c** was analyzed using a one-way analysis of variance (ANOVA). Data in **f-h** were analyzed using a two-way ANOVA followed by a Tukey’s multiple comparison test with 95% confidence interval. Data in **i** was analyzed using a one-way ANOVA followed by a Dunnett’s multiple comparison test with 95% confidence interval. *, ** and **** indicate P<0.05, P<0.01, and P<0.0001, respectively; N.S., not significant.

Work by others has demonstrated that activated astrocytes become significantly smaller compared to quiescent astrocytes *in vivo*.^12^ The aberrant morphologies coincide with high expression of *gfap* during activation.^52^ In our brain hydrogel, astrocytes retain a predominantly stellate morphology in both standard and quiescent medium, and shift to a mix of rounded, polarized, and stellate cells in cytokine medium (Fig. 5d-e). In collagen, astrocytes exhibited a heterogeneous range of morphologies as expected in activated populations (Fig. 5d-e).^52^ Similar to A1 activated astrocytes described by Liddelow *et al*.,^12^ we found astrocyte process length, volume, and surface area decreased significantly in the highest cytokine concentration in the brain hydrogel, while there was no significant difference across collagen cultured conditions (Fig. 5f-h). Astrocytes have also been shown to become highly migratory upon activation *in vivo*^*5,53*^. We expectedly found highly migratory astrocytes on 2D TCPS^54^, and that cells migrated significantly more in brain hydrogels with cytokine medium, compared to the standard culture in the control brain hydrogel (Fig. 5i).

HA is a polysaccharide in brain tissue that has been commonly incorporated into *in vitro* ECM models of the brain^25,43,55^. Others have shown correlation between concentration of low MW HA in a biomaterial and astrocyte activation^10,21^, and disease progression (e.g. glioblastoma).^21,43,55,56^ We also verified that low MW HA induced astrocyte activation in our brain hydrogel. We incorporated thiolated low MW HA (10kDa) in the brain hydrogel and stained for GFAP after 48 hours (Fig. S10). We found that GFAP expression increased in a dose-dependent manner, with 0.1mM HA being the lowest dose at which astrocytes remained quiescent (Fig. S10), in agreement with reports by others in glial cell studies.^10,21,56^ Together these data demonstrate that astrocytes cultured in the brain hydrogel remain quiescent *in vitro* and can undergo cytokine- and HA-mediated activation as defined by morphology, expression of GFAP and vimentin, and a migratory phenotype. Further, quiescent astrocyte populations were observed in the synthetic brain-mimicking hydrogel, and not in collagen.

## Outlook

Bioengineers have designed *in vitro* models that can recapitulate the brain ECM^6-10,22^. However, there is no tunable *in vitro* model that can maintain astrocyte quiescence. The instructive design of this brain ECM specific hydrogel allows for tunable control of astrocyte activation, in contrast to protein- and sugar-based models like collagen and hyaluronic acid. An exciting recent finding illustrated how proteins secreted by cells within synthetic hydrogels regulate cell phenotype^57^, potentially putting into question the need for inclusion of ECM binding and degradation sites into gels. However, we found here that these initial signals are very important in driving astrocyte quiescence and activation. For example, astrocytes encapsulated in hydrogels with *either* integrin-binding *or* MMP-degradable peptides did not show a stellate morphology (Fig. 3). The need for these tissue-specific hydrogels is of particular importance to instruct cells that do not produce their own matrix, for example oligodendrocytes and schwann cells^58^, and it has been shown many times that these cues can impact cell differentiation^58,59^ and viability^39^. Similarly, specific ECM cues provided by integrin-binding sites^8,28,60^, inflammatory molecules^12,61^, and mechanical properties^40^, have been shown to influence the deposition of ECM proteins by astrocytes. We foresee the use of this synthetic brain hydrogel to understand extracellular mechanisms of astrocyte activation, as well as individual contributions of quiescent and reactive astrocytes with other CNS cells in neurodegenerative diseases. In fact, recent studies demonstrate the need to control for and study the dynamic cell-to-cell interactions, especially in the context of neurons, astrocytes, and microglia^62,63^ to better study Alzheimer’s disease *in vitro*. Rational biomaterial design is also critical for to accelerate and/or support proper brain tissue repair after stroke ^64^. Future studies investigating the functional interactions amongst these various neural cell types in our tunable system will allow for precise and controlled manipulation, experimental schemes to tease apart the complex signaling mechanisms underlying cell-to-cell communication in the context of neurodevelopmental and neurodegenerative diseases, and potentially offer synergistic therapeutic value.

## Acknowledgements

The authors would like to thank Sarah Perry for providing peptide synthesizer equipment, the NIH Neurobiobank for providing human brain tissue, and support from Lauren Jansen on hydrogel design workflow. We thank Dr. Stephen Eyles and the Mass Spectrometry Core Facility at the Institute of Applied Life Sciences for support with LC-MS/MS. This work was supported by NIH Supplement Award 3DP2CA186573-01S1 to SG, NIH New Innovator Award to SRP (1DP2CA186573-01), an NSF CAREER to SRP (DMR1454806), a grant from the ONR N00014-17-1-2056 to AJC and SRP, a Northeast Alliance for the Graduate Education and Professoriate (NEAGEP) fellowship to SG, and a Spaulding-Smith Fellowship to SG. SRP is a Biomedical Scholar funded by the Pew Foundation.

## Author contributions

S.G. contributed to drafting the manuscript, data collection, and data interpretation. AJC and CP contributed to data analysis and interpretation, and editing the manuscript. S.R.P. contributed to conceptual design, data interpretation, and drafting the manuscript.

## Methods

### Cell Culture

Normal human astrocytes of cortical origin were purchased from ScienCell Research Laboratories (San Diego, CA USA) and maintained in Astrocyte medium (ScienCell, San Diego, CA USA). Primary cells were used between passages 1 to 3 for all experiments. MDA-MB-231-BrM2a cells were a generous give from Joan Massague, and MDA-MB-231 cells were a generous gift from Dr. Sallie Schnedier lines. Both 231 cell lines were cultured in DMEM with 10% fetal bovine serum (FBS, Thermo). The Neuro2A, mHippoE18, and SK-N-AS were cultured in DMEM supplemented with 10% FBS, 0.1mM NEAA, 1% L-glutamine, and 1mM sodium pyruvate.

### Acquisition of human brain tissue

Deidentified human brain tissues were obtained from the NIH NeuroBioBank at Harvard Medical School. All tissues had been flash-frozen from four post-mortem neurologically normal individuals: 3 females, and one male from the ages of 30-60.

### Mass Spectrometry

The CNMCS Compartment Protein Extraction Kit (Millipore, Sigma) was used to decellularize all human brain tissue samples as described in^29^. Briefly, intracellular soluble proteins were extracted with sequential incubation in the appropriate buffers according to manufacturer’s instructions. This resulted in an insoluble ECM-rich pellet. The ECM-rich pellet of four brain tissue samples was solubilized and reduced in 8 M Urea, 100 mM ammonium bicarbonate, and 10 mM dithiothreitol (DTT, Thermo Fisher Scientific, Waltham, MA) for 30 minutes at pH 8 and 37°C. Samples were then alkylated with 25mM iodoacetamide (Sigma-Aldrich) in the dark at RT for 30 minutes. Samples were subsequently quenched with 5mM DTT and the solution was diluted to 2M urea with 100 mM ammonium bicarbonate (pH 8). Proteins were digested via trypsin (Thermo) and Lys-C endoproteinase (Promega, Madison, WI) at a weight ratio of 1:50 enzyme to protein overnight (12-16 hours) at 37°C. Finally, samples were cleaned and concentrated using a C18 column (Thermo).

Digested peptides were separated by reverse phase LC gradient prior to mass spectrometry analysis with an Orbitrap Fusion Tribid (Thermo). Identified peptides were aligned against the Matrisome using the Thermo Proteome Discoverer 1.41.x. Parameters for analysis used trypsin as a protease, with 4 missed cleavages per peptide, a precursor mass tolerance of 10 ppm, and fragment tolerance of 0.6 Da.

### Determination of Young’s modulus for brain tissue and hydrogels

The Young’s modulus was measured using an indentation custom built instrument used as previously described.^45^ Briefly, a flat cylindrical punch of mm diameter was brought into contact with the hydrogel at a fixed displacement rate of 15 *μ*m/s, for a maximum load of 1.5mN. The linear response of the low strain regime was analyzed using a Hertzian model accounting for dimensional confinement between the contact radius (a) and the sample height (h) (0.5 < a/h <2) as described previously^65^. To tune the hydrogel Young’s modulus to match the brain, PEG-Maleimide hydrogels were formed at different macromer concentrations containing the integrin-binding and MMP-degradable peptides and swollen in PBS buffer overnight. Finally, the Young’s modulus was similarly obtained for six individual porcine brains (n>=5) and 3 human brain donors (n=2). All samples had a thickness of 1mm to ensure the dimensional confinement remained within 0.5 < a/h < 2.

### Solid-phase peptide synthesis

Peptides (GRGDSPCG, GCALMKYHILNTLQCSE, GCDPGIVRRADRAAVP, GCDPGIKVAV, GCDPGYISGR, GCGDGEA, GCGFYFDLR, CSVTCG, CGGAEIDGIEL, GCRDIPVSLRSGDRCG, GCRDRPFSMIMGDRCG, GCRDVPLSLTMGDRCG, GCRDVPLSLYSGDRCG, and GCRDIPESLRAGDRCG) were synthesized on a CEM Liberty Blue automated solid phase peptide synthesizer (CEM, Mathews, NC) using FMOC protected amino acids (Iris Biotech GMBH, Germany). The peptide was cleaved from the resin by sparging-nitrogen gas through a solution of trifluoroacetic acid (TFA), triisopropylsilane (TIPS), 2,2’(Ethylenedioxy)diethanethiol (DODT) and water at a ratio of 92.5:2.5:2.5:2.5 % by volume, respectively (Sigma-Aldrich) for 2-3 hours at RT in a reactor vessel (ChemGlass, Vineland, NJ). After reaction, the solution was filtered, and the peptide was precipitated using ethyl ether at −80°C (Thermo). The molecular weight of the peptide was validated using a MicroFlex MALDI-TOF (Bruker, Billerica, MA) using alpha-cyano-4-hydroxycinnamic acid as the matrix (Sigma-Aldrich). Peptides were analyzed and purified to >95% on a VYDAC reversed-phase C18 column attached to a Waters 2487 dual (lambda) adsorbable detector and 1525 binary HPLC pump (Waters, Milford, MA).

### Cell adhesion assay to brain ECM peptide surfaces

Integrin-binding peptides were attached to 15mm glass coverslips by adapting a previous published approach.^66^ Coverslips were treated with UV/ozone at 1 atm (UV/Ozone ProCleaner, 180 *μ*g/m^3^ ozone level in chamber, 4.6 mW/cm^2^ peak UV intensity, 10.15W UV lamp power requirement, BioForce Nanosciences, Salt Lake City, UT) for 10 minutes to expose -OH groups. (3-aminopropyl)triethoxysilane (APTES, Sigma-Aldrich, St. Louis, MO) was reacted with the glass surfaces in a 90°C oven via vapor deposition overnight, wrapped in foil. The glass coverslips were sequentially rinsed three times in Toluene (Thermo), 95% ethanol (Thermo), and distilled water. The glass was allowed to dry in the oven at 90°C for 1 hour, and then functionalized with 10 g/L N,N-disuccinimidyl carbonate (DSC, Sigma-Aldrich) and 5 v% N,N-diisopropylethylamine (DIEA, Sigma-Aldrich) in acetone (Thermo), sequentially, for 2 hours each. Coverslips were then rinsed with acetone, air-dried for 10 minutes, and either used immediately or stored in a desiccator overnight.

Each glass coverslips surface was coated with 70µL of integrin-binding peptide solution, allowed to react for 2 hours, rinsed 3 times with pH 7.4 PBS, and then blocked with 10 µg/cm^2^ MA(PEG)_24_ (Thermo) for 2 hours to prevent nonspecific protein adsorption. The coverslips were glued to the surfaces of each well in a 24-well plate with epoxy (Thermo), rinsed 3 times with PBS, and UV sterilized for 1 hour before cell seeding. Cells were seeded at 10,000 cells per cm^2^ to functionalized glass coverslip surfaces in serum free DMEM and imaged by a controlled Zeiss Axio Observer Z1 microscope (Carl Zeiss, Oberkochen, Germany) using an AxioCam MRm camera and an EC Plan-Neofluar 20X 04 NA air objective. Images were taken every five-minutes for an incubation period of 1-, 2- and 12-hours. Cell area was quantified by manual tracing in ImageJ (NIH, Bethesda, MD).

### Competitive binding assay

Cells were seeded at 10,000 cells per cm^2^ onto coverslips functionalized with integrin-binding peptides after 30 minutes of pretreatment with individual peptides or the complete brain-ECM peptide mixture. Cells were seeded at 10,000 cells per cm^2^ onto glass coverslips (15mm, Thermo) functionalized with the individual peptides or a 500 ug/mL (35 ug/cm^2) complete brain-ECM peptide mixture: 35wt% RGD, 8wt% LRE, 5wt% IKVAV, 2wt% PHSRN-RGDS, 13wt% DGEA, 10wt% FYFDLR, 4wt% VTCG, 1wt% AEIDGIEL, 3wt% ALMKYHILNTLQCSE, 3wt% IVRRADRAAVP, 5wt% TWSKVGGHLRPGIVQSG, 2wt% GRKRK, 7wt% YIGSR, and 1wt% GWTVFQKRLDGS. Conditions with individual peptides were at the same concentration as the peptide is represented in the full mixture. Cells were imaged beginning at 5 minutes after seeding in an environment-controlled Zeiss Axio Observer Z1 microscope. Images were taken at five-minute intervals for 2 hours, or every fifteen-minutes for 12 hours, and cell areas were manually traced in ImageJ.

### Synthesis of 3D Synthetic Brain Hydrogels

A 20K 4-arm PEG-maleimide (Jenkem Technology, Plano, TX) was reacted with 4mM integrin-binding peptides for 5 minutes in serum-free medium at pH 7.4 (solution A). The crosslinker solution (solution B) was composed of 77 molar % of 1K linear PEG-dithiol (Jenkem) and 13 molar % of the MMP-degradable peptide cocktail. These two solutions were combined at a 1:1 molar ratio of thiol to Maleimide in TEA at pH 7.4 for a total final gel volume of 10uL. For experiments with cells, cells were added to “solution A” by using solution A as the fluid to resuspend a cell pellet prior to gelation with “solution B”. Gels were allowed to polymerize for 10 minutes prior to addition of cell culture medium.

To ensure that the brain hydrogels stayed in a fixed location during imaging, we prepared coverslips that would covalently crosslink to the gels during network polymerization, as previously described^67^. Glass coverslips were UV/ozone treated for 10 minutes and functionalized with 2 vol% solution of 3-mercaptopropyl-trimethoxysilane (MPT, Thermo) in 95% ethanol (adjusted to pH 5.0 with glacial acetic acid) for 1 hour. The wells were rinsed 3 times with 100% ethanol and allowed to air dry for 10 minutes before addition of the brain hydrogel solutions.

### Validation of peptide incorporation

The Measure-iT thiol kit was used to quantify unreacted thiols (Thermo) as previously described.^39^ Integrin-binding peptides were incorporated into gels at 2 and 4mM concentrations in a 100 µL volume of PEG-maleimide for 10 minutes before reacting hydrogels with 100µL of the Measure-iT kit working solution. Separately, dithiol terminated crosslinkers (PDT at an equimolar ratio of thiol to maleimide, or MMP-degradable peptide (13mole%) with 87mole% PDT) were reacted with the PEG-maleimide in 10*μ*l aliquots for 10 minutes before reacting with 100µL of the kit working solution. These hydrogels were incubated in sodium borohydride (NaBH4, Sigma) in water at a molar ratio of 4:1 NaBH4 to thiol for 4 hours before adding the thiol kit working solution. Hydrogel supernatant and solutions were acquired and read at an excitation of 494nm and emission of 517nm, as we have previously reported ^68^. To detect any peptides that did not incorporate into the hydrogel network, we removed the supernatant and analyzed it using a MicroFlex MALDI-TOF (Bruker) with alpha-cyano-4-hydroxy cinnamic acid (Sigma) as the matrix.

### Human Astrocytes spreading and variation of integrin-binding peptide concentration

Human astrocytes were encapsulated into the brain customized hydrogel with integrin-binding peptide concentrations varying from 0 to 4 mM at a density of 5,000 cells/µL. After 24 hours, hydrogels were fixed with 4% formaldehyde (Acros) for 10 minutes and stained with GFAP (Abcam ab7260), CellMask Membrane stain (Thermo C10046) and DAPI (Sigma). Cells were imaged on a Zeiss Spinning Disc microscope (Zeiss) using an HRm AxioCam. Images were processed with Zen software (Zeiss) and cell areas were manually traced with ImageJ (NIH).

### Collagen Hydrogel Synthesis

Type I collagen (rat tail, Corning, NY, lot 7079004) were formed by combining with NaOH, 10X DMEM (Sigma) and cells in cold, serum-free medium. Gels were allowed to polymerize at 37°C for 30 minutes before adding cell culture medium.

### Control, Quiescent, and cytokine-containing astrocyte media

Serum-free base medium containing 50% neurobasal (Thermo), 50% DMEM, 100 U/ml penicillin, 100 ug/ml streptomycin, 1 mM sodium pyruvate (Thermo), 292 ug/ml l-glutamine (Thermo), and 5 ug/ml of N-acetyl cysteine (Sigma) were supplemented with heparin-binding epidermal growth factor (HBEGF, Peprotech, Rocky Hill, NJ Cat #100-47) at 5 ng/ml as previously described^12^. 1X SATO was incorporated from 100X SATO aliquots (10 mg/ml BSA (Thermo), 10mg/ml Transferrin (Sigma), 1.6 mg/ml Putrescine (Sigma), 6 ug/mL progesterone (Sigma), and 4ug/ml sodium selenite (Sigma)) at the moment of the experiment.

Astrocytes were cultured in either this quiescent medium or control human astrocyte medium from ScienCell (ScienCell, San Diego, CA USA). For dosing with cytokines, quiescent medium was treated with IL-1*α* (3 ng/mL, Sigma, #I3901), TNF-*α* (30 ng/ml, Cell Signaling Technology, Danvers, MA #8902SF) and C1q (400 ng/ml, MyBioSource, San Diego, CA #MBS143105) for the single dose cytokine conditions (+). Double cytokine conditions (++) were dosed with double the cytokine concentrations listed above.

Passage 1-3 primary human astrocytes were cultured in the brain hydrogel for 24 hours in either control or quiescent astrocyte medium. After 24 hours, the medium was changed to that of control, quiescent, single dose- or double dose-cytokine medium. Following 24 hours after cytokine dosing, the medium was removed, and hydrogels were fixed in 4% PFA for 10 minutes at 37°C before immunocytochemistry.

### Astrocyte culture in synthetic brain hydrogels containing ultra-low MW HA

To functionalize the brain hydrogel with low MW hyaluronate thiol (MW 10kDa, Creative PEG Works, HA-371), HA containing thiol groups was reconstituted in PBS and incorporated with the crosslinker mixture composed of 77 molar % of 1K linear PEG-dithiol (Jenkem) and 13 molar % of the MMP-degradable peptide cocktail where HA was added to a final concentration of 0.1-, 0.5- and 1mM HA in a 10 uL hydrogel. Astrocytes were encapsulated in these HA-containing brain hydrogel as described earlier. Hydrogel contained 4mM integrin-binding peptides and were crosslinked at a 1:1 molar ratio of thiol to Maleimide in TEA at pH 7.4. Gels were allowed to polymerize in 10uL volumes with 5,000 cells/**μ**l, and cell culture media was added after 10 minutes. Astrocytes were cultured for 24, 48, and 72 hours in the HA containing hydrogels where medium was removed and replaced with 4% PFA to prepare the hydrogels for immunocytochemistry.

### Immunocytochemistry

Hydrogels were fixed in 4% paraformaldehyde solution for 10 minutes at room temperature, then washed 3X with ice-cold PBS. Gel-embedded cells were blocked with 5% Bovine Serum albumin (BSA, Thermo) for 30 minutes, followed by incubation with the CellMask membrane stain (Thermo C10046) for 1 hour in the dark at RT, then washed 3X with PBS. To quantify activation, gel-embedded cells were permeabilized with 0.25% Triton X-100 (Sigma) for 10minutes, rinsed with 0.1% Triton X-100 for 3 minutes, washed with PBS 3X over 5minutes, blocked with 5% BSA at room temperature for 30 minutes, and incubated with primary antibodies to GFAP (1:1000 ab7260, Abcam), vimentin (1:200 ab24525, Abcam) and Cell Mask membrane stain (1:1000), overnight at 4°C. Finally, gels were washed and incubated in secondary antibody solutions (1:500 ab96883, Abcam, Goat anti-rabbit, green and 1:500 ab96950, Abcam, Goat anti-chicken, red, respectively) for 1 hour at RT. Cells were imaged on a Zeiss Spinning Disc confocol (Zeiss) using an HRm AxioCam. Images were taken using Zen software (Zeiss) and cell morphology was analyzed in ImageJ (NIH) and Imaris (BitPlane, Belfast, UK). Normalized GFAP expression was determined quantitatively from the ratio of the average fluorescence per pixel in the cell body and processes to the average intensity of the background as quantified with ImageJ (NIH). The number of GFAP-positive cells were counted using the 3D cell counter plug-in in ImageJ.

Three dimensional reconstructions and filament traces of the astrocytes were generated from confocal Z-stacks of the CellMask membrane stain using Imaris (Bitplane) and ImageJ (NIH) to obtain quantitative morphological data. Sholl^69^ analysis was performed to quantify astrocyte process number and length. Total additive process length, degree of branching (# process ends / # primary processes, and cell diameter were calculated for each cell.

### Statistical Analysis

Statistical analysis was performed with GraphPad Prism (7.0d) (GraphPad Software, Inc., La Jolla, CA). Statistical significance was evaluated using a one-way analysis of variance (ANOVA) followed by a Tukey’s post-test for pairwise comparisons. P-values <0.05 are considered significant, where p < 0.05 is denoted with *, < 0.01 with **, < 0.001 with ***, and < 0.0001 with ****.

