## Supplemental Materials for "Control of Astrocyte Quiescence and Activation in a Synthetic Brain Hydrogel"

**Supplementary Figures**

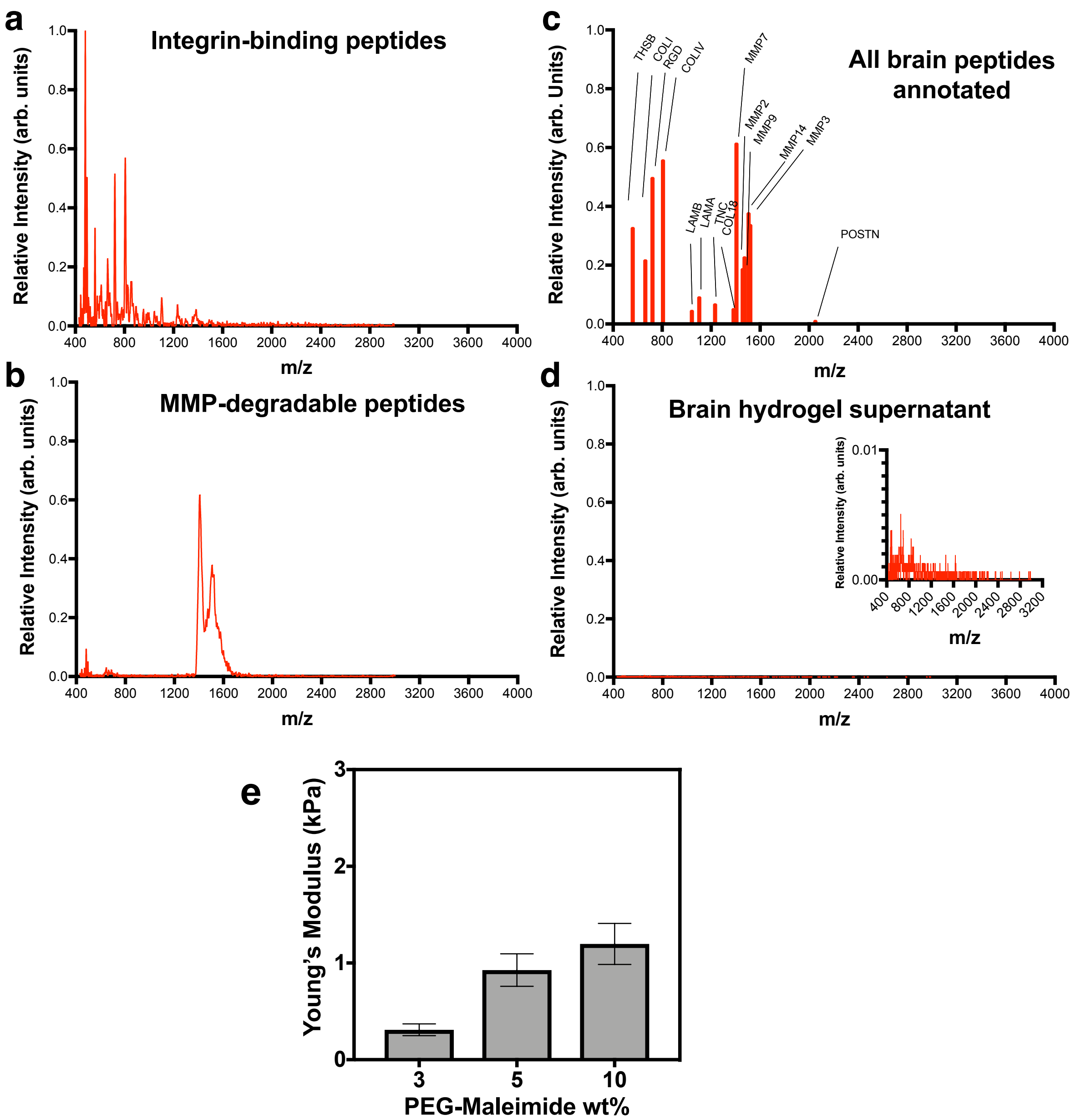


**Figure S1.** Characterization of brain hydrogel peptide incorporation and modulus. **a**) Spectra of adhesive and **b**) degradable peptides as identified via MALDI-TOF analysis. **c**) Peaks corresponding to the different amino acid sequences in the mixture. **d**) Analysis of brain hydrogel supernatant after polymerization was performed for 10 minutes shows no peaks. **e)** Modulus of hydrogels as a function of the weight % of PEG-maleimide included.

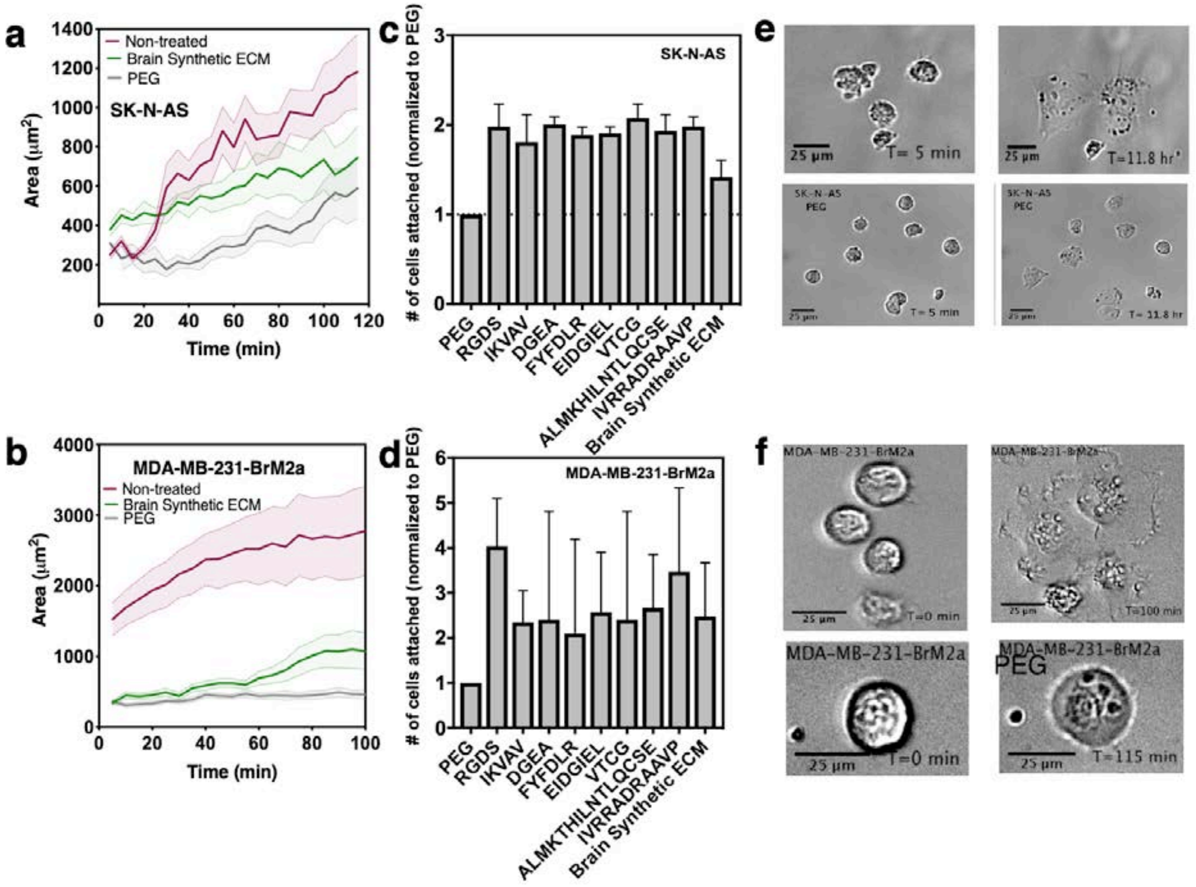


**Figure S2.** Adhesion of human cell lines to integrin-binding peptides in the brain hydrogel formulation. Human cell lines **a**) SK-N-AS (neuroblast) and **b**) MDA-MB-231-BrM2a (brain-metastatic breast cancer) were seeded in surfaces functionalized with poly(ethylene)glycol (PEG) (gray), brain integrin-binding peptides (maroon), or previously incubated with the brain integrin-binding peptide cocktail (green) and then seeded into a coverslip functionalized with the same peptide mixture. Cell area was traced every five minutes for a period of 120 (a) or 100 minutes (b). Cell area increased significantly in the non-treated (maroon) cells that were seeded directly onto coverslips with the adhesive peptide mixture, while the cell area remained small in the PEG condition. Cells treated with the adhesive peptide mixture for five minutes prior to seeding onto coverslips took longer to increase the cell area. Data shown is mean (solid line) + s.d. (shade region). N=2, *n=30.* **c**) SK-N-AS and **d**) MDA-MB-231-BrM2a cells were seeded into coverslips functionalized with each integrin-binding peptide for a period of 2 hours. Cells were washed, fixed and stained with DAPI and counted with ImageJ software. Data shown is the number of cells attached normalized to PEG as mean + s.d. N=2, *n=3.* **e**) Representative images of SK-N-AS and **f**) MDA-MB-231-BrM2a cells adhering overtime to coverslips functionalized with the integrin-binding peptide mixture or PEG. Cells show a larger area when cultured on coverslips functionalized with the adhesive peptide mixture in contrast to PEG alone.

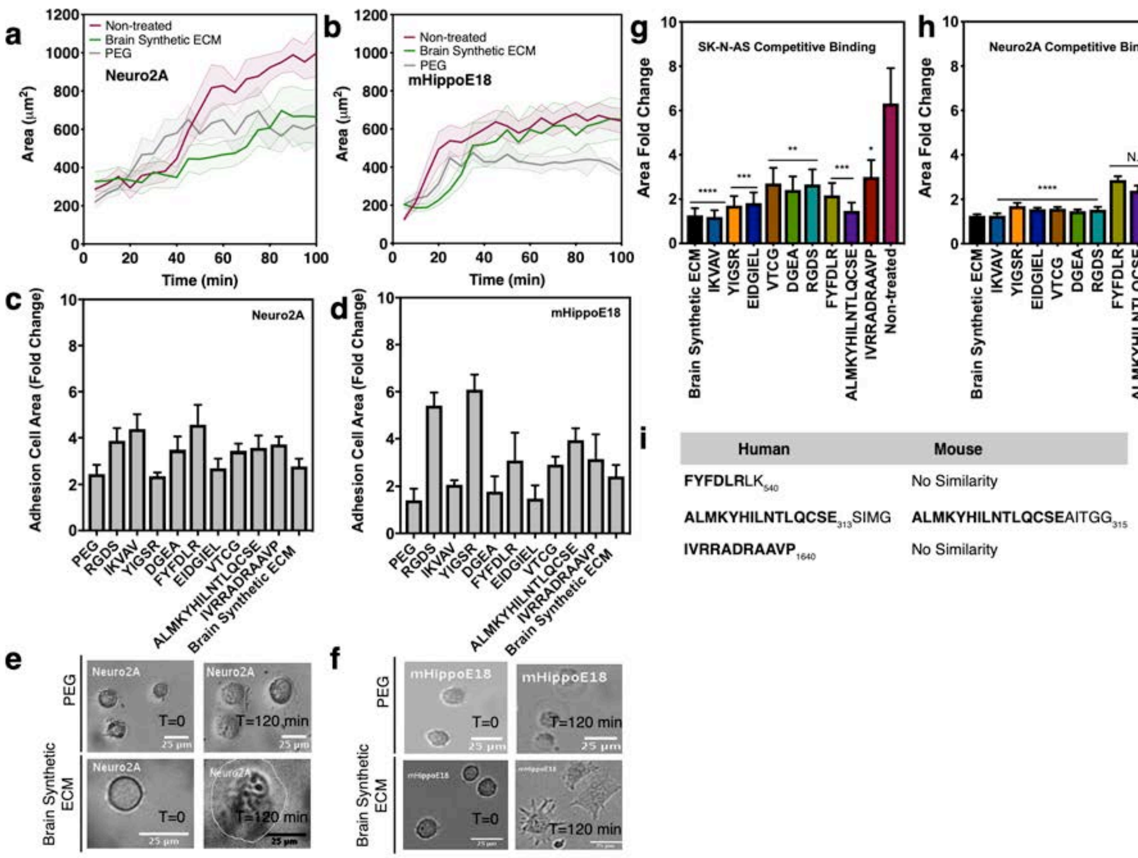


**Figure S3.** Analysis of mouse cell adhesion to human brain peptides and comparative sequence homology. **a**) Quantification of cell area traces overtime for the murine cell lines Neuro2A and **b**) mHippoE18. Cells were seeded onto coverslips functionalized with poly(ethylene)glycol (PEG) (gray), the brain integrin-binding ECM peptides (maroon), or incubated for five minutes with the integrin-binding brain peptides followed by seeding onto coverslips functionalized with the same integrin-binding peptides (green). Data shown is mean (solid line) +/- s.d. (shaded area). N=2, *n=30.* **c**) Cell lines Neuro2A (mouse neuroblastoma) and **d**) mHippoE18 (mouse embryonic hippocampus) were seeded into coverslips functionalized with each integrin-binding peptide and allowed to adhere for 2 hours. Change in cell area from directly after seeding up to 2 hours are shown for each integrin-binding peptide, PEG, and the brain integrin-binding peptide mixture. **e**) Neuro2A and **f**) mHippoE18 representative images of cell spreading on coverslips functionalized with PEG or the brain integrin-binding ECM mixture over 2 hours. **g**) Competitive binding of the human cell line SK-N-AS and **h**) murine cell line Neuro2A. Cells were previously incubated for five minutes in the peptide of interest in soluble form and subsequently seeded into a coverslip functionalized with the integrin-binding peptide mixture. Data shown is mean + s.e.m. N=2, *n=30.* **i**) Sequences that did not inhibit cell adhesion of the Neuro2A cell line to the integrin-binding peptide mixture functionalized coverslip are not found in the protein sequence of the mouse.

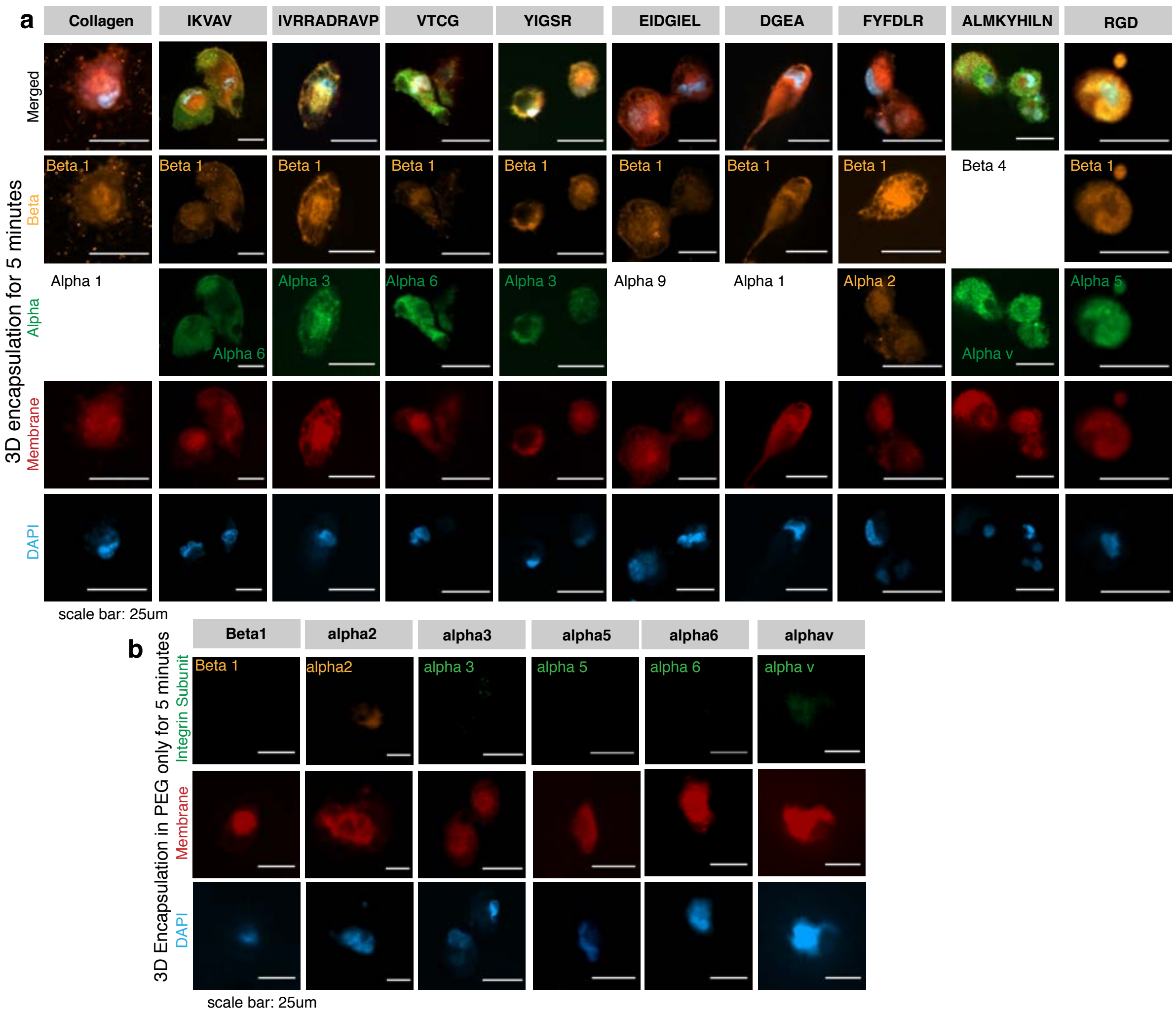


**Figure S4.** Validation of integrin localization at cell membrane after encapsulation. **a**) Human primary astrocytes were encapsulated for five minutes in collagen or hydrogels modified with a single integrin-binding peptide at the same concentration as in the brain adhesive peptide mixture, **b**) or negative control poly(ethylene)glycol hydrogels without integrin-binding peptides. Hydrogels were fixed and stained for the expected integrin subunits (colors in the text inset for each integrin subunit) known to bind with their heterodimer partners to the peptides in the gel (See Tables S2-3 for full list). Membrane stain (red) and DAPI (blue) were also included for visualization purposes. Omitted images are from unreliable antibodies found for human integrin subunits (not suitable for immunofluorescence).

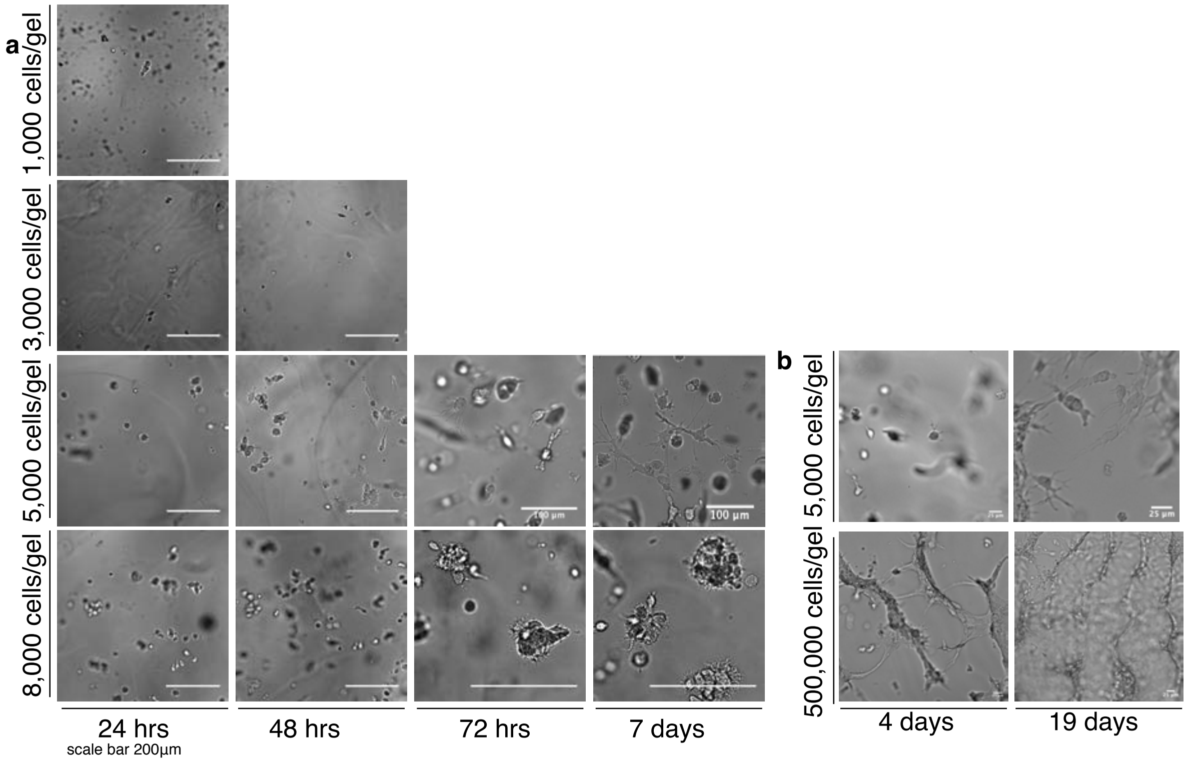


**Figure S5.** Optimization of human primary astrocytes seeding density in the brain hydrogel. **a**) Human primary astrocytes were encapsulated at a density of 1,000-, 3,000-, 5,000-, and 8,000 cells/gel (10uL) in the brain hydrogel and cultured for a period of 7 days. **b**) Cells can also be encapsulated at greater densities (e.g. 500,000 cell/gel), which results in undesired astrocyte network formation.

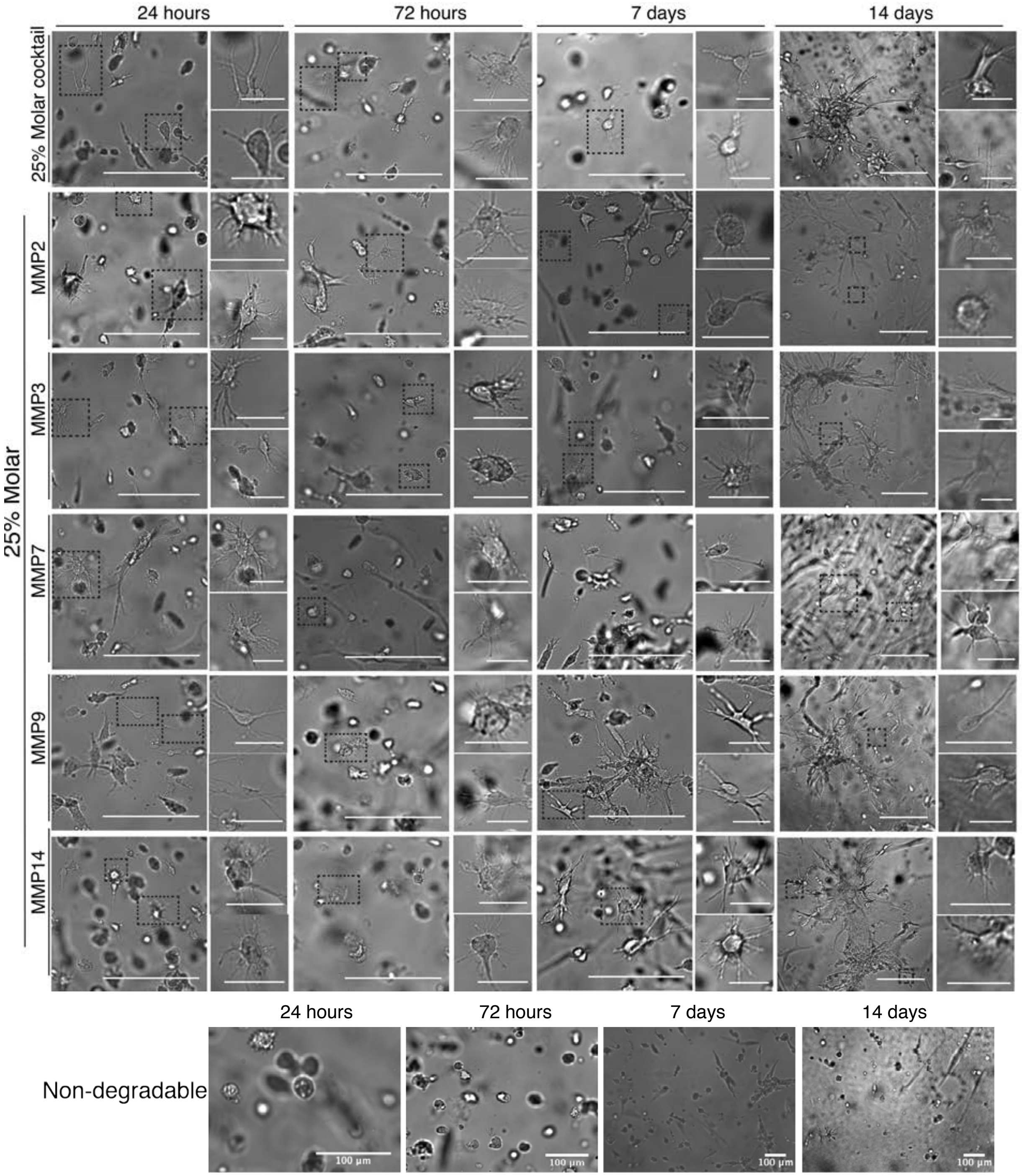


**Figure S6.** Human primary astrocytes extend processes through networks with each MMP-degradable domain. Representative images of human astrocytes encapsulated in a functionalized brain hydrogel, hydrogels with a single MMP-degradable peptide and an integrin-binding peptide concentration of 2mM, or a non-degradable PEG hydrogel for a period of 14 days. Scale bars for large images = 200um, and 50um for insets.

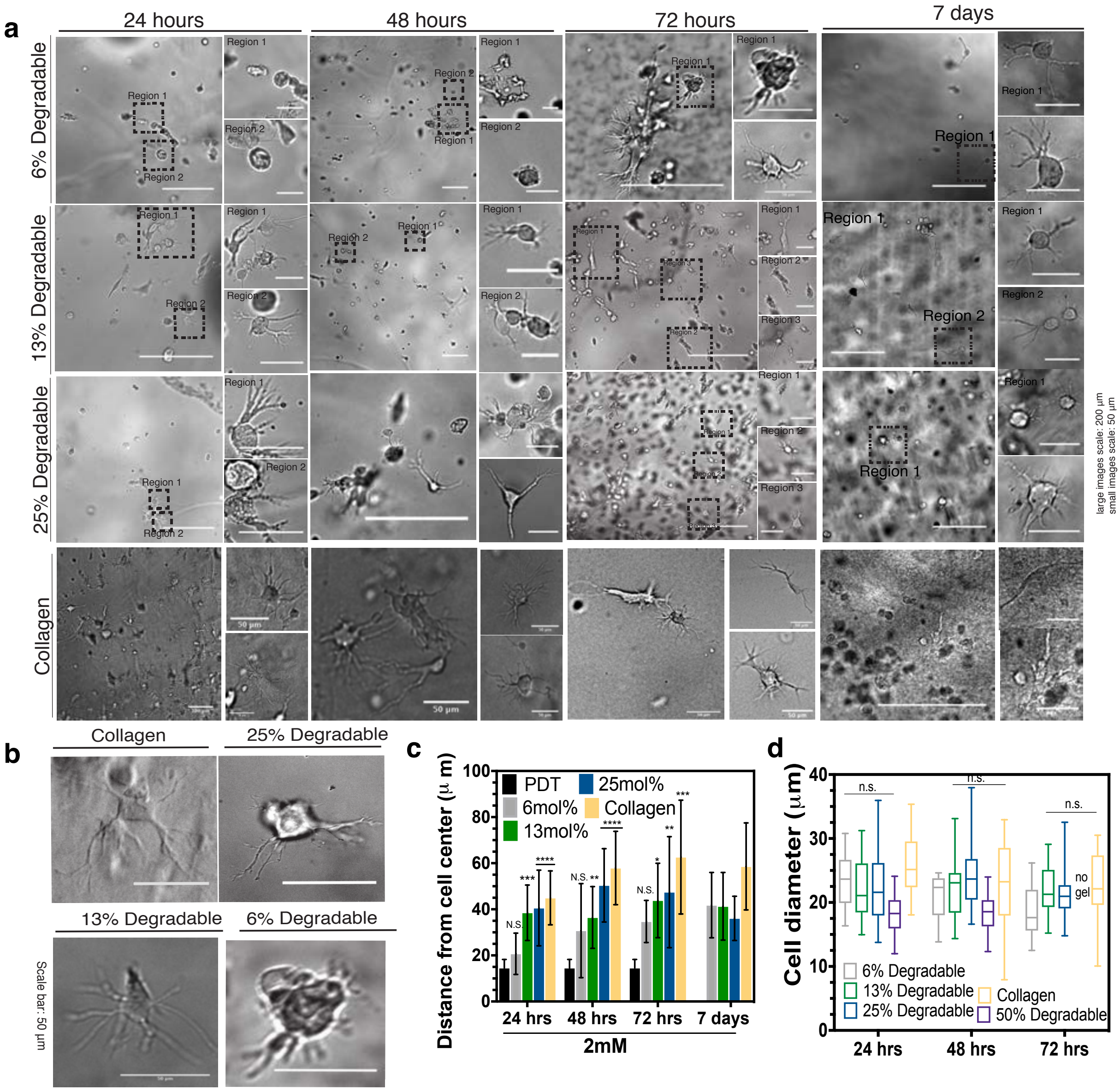


**Figure S7.** Optimization of MMP-degradable peptide concentration in the brain hydrogel. **a**) Human astrocytes were encapsulated in collagen and brain hydrogels functionalized with 6mol%, 13mol%, and 25mol% MMP-degradable peptide mixtures at a constant 2mM integrin-binding peptide concentration. Representative images of cell morphology and dynamic remodeling of the hydrogel for a period of 7 days. **b**) Representative images of astrocyte cell morphology after 72 hours of encapsulation. **c**) Quantification of the distance from the cell center over time for hydrogels at different conditions as compared to a non-degradable PEG hydrogel. Astrocytes encapsulated in a brain hydrogel with 13mol% and 25mol% MMP-specific peptide mixture achieve a longer extension similar to collagen cultured astrocytes. **d**) Cell diameter of astrocytes cultured in brain hydrogels at different conditions, and collagen for 72 hours. Data shown is mean + s.d. N=3, *n=4.*

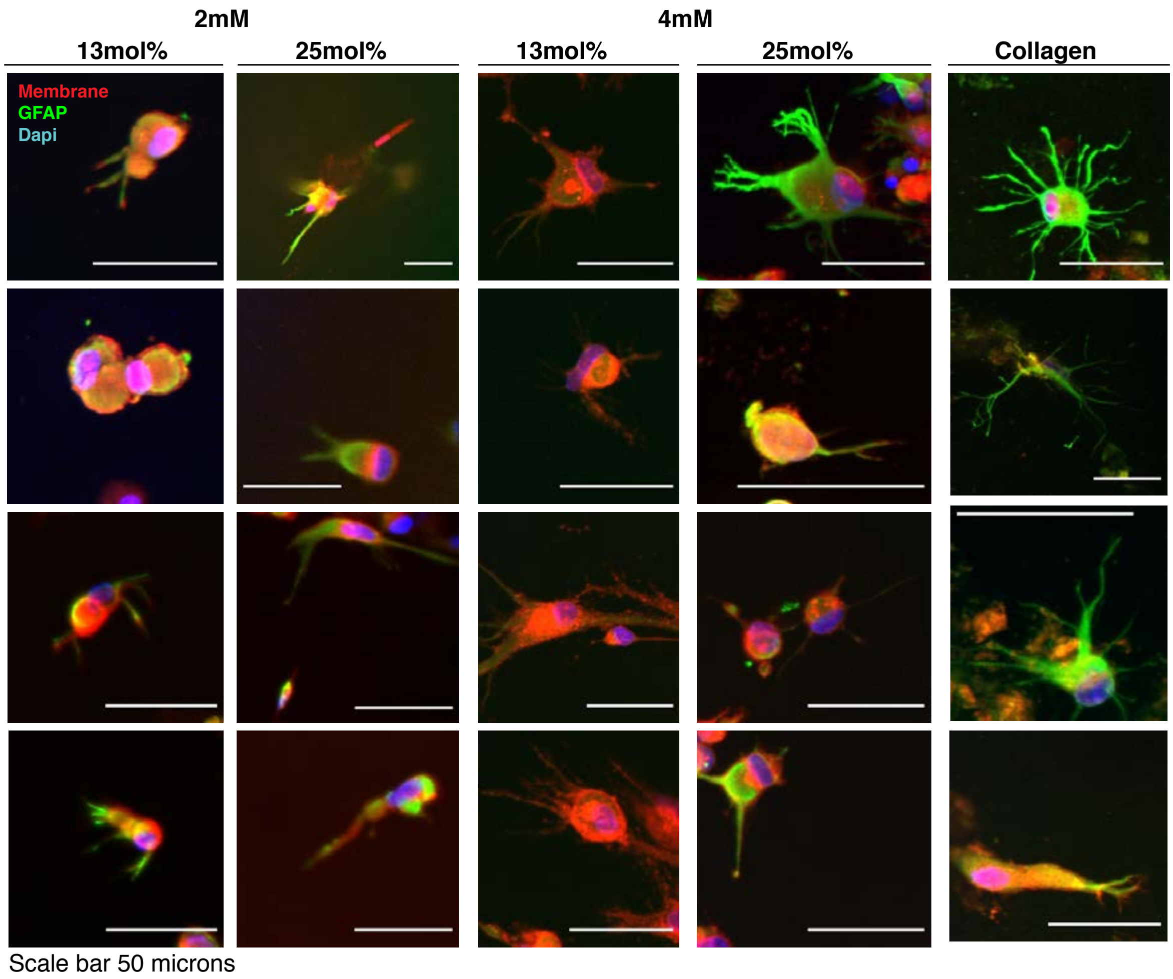


**Figure S8.** Astrocyte activation is regulated by tuning of integrin binding and MMP-degradable peptides in the brain hydrogel. Representative images of astrocytes stained for *gfap* (green), membrane (red) and DAPI (blue) at different formulations of the brain hydrogel as compared to collagen. Concentration of integrin-binding peptides is shown in [mM] with the MMP-degradable peptide concentrations in mole%.

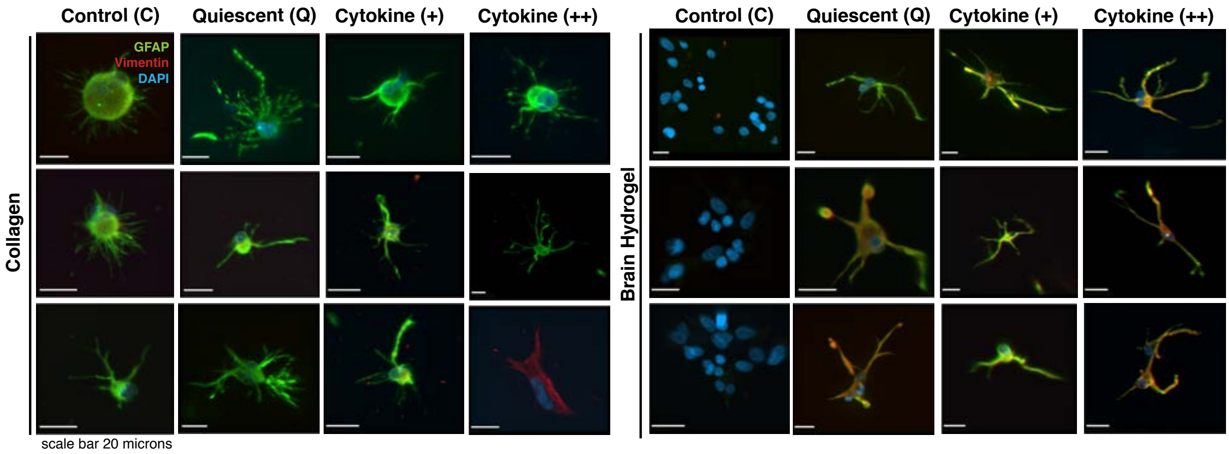


**Figure S9.** Astrocytes are activated with cytokine dosing in the brain hydrogel. Representative images of astrocytes encapsulated in collagen or brain hydrogel in the presence of control medium (C), quiescent medium (Q), or cytokine-dosed conditions with either a normal concentration (+), or double (++).

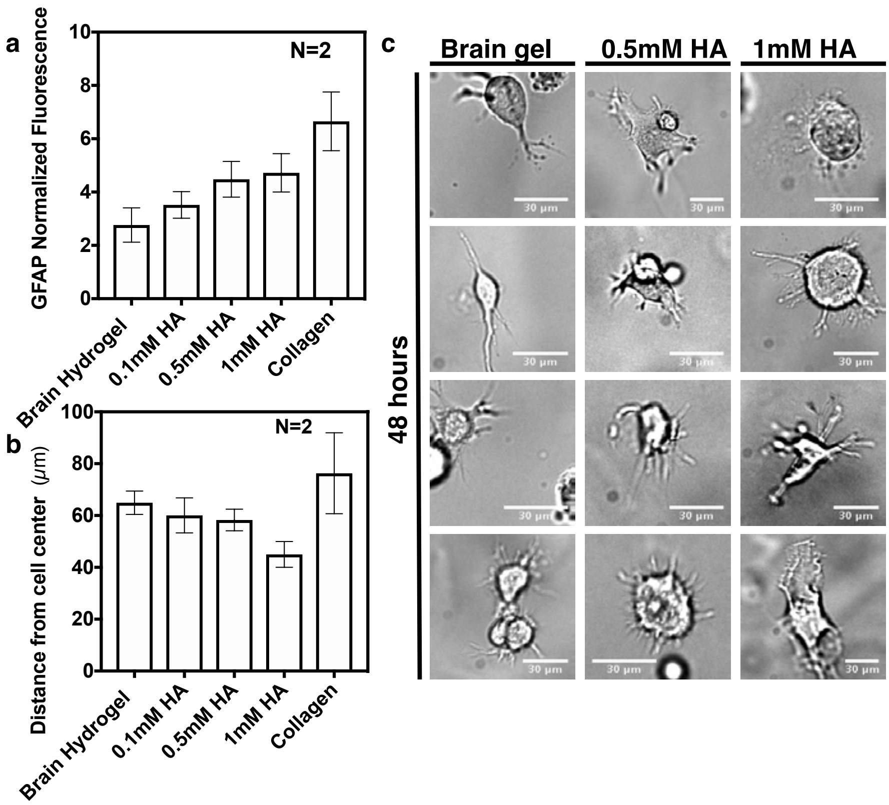


**Figure S10.** Human primary astrocytes become activated to small molecular weight hyaluronic acid in a dose-dependent manner. **a**) Normalized GFAP fluorescence increases with increasing concentration of hyaluronate thiol in the brain hydrogel. Data shown are mean and s.e.m. N=2, *n=3.* **b**) Distance from the cell center for astrocytes encapsulated in hyaluronate thiol modified brain hydrogels for 48 hours. Process length decreases with increasing concentration of HA. Data shown are mean and s.e.m. N=2, *n=3.* **c**) Representative bright field images of astrocytes encapsulated in hyaluronate thiol modified brain hydrogels for a period of 48 hours. Scale bars = 30um.
